# Objective Evaluation of Multiple Sclerosis Lesion Segmentation using a Data Management and Processing Infrastructure

**DOI:** 10.1101/367557

**Authors:** Olivier Commowick, Audrey Istace, Michaël Kain, Baptiste Laurent, Florent Leray, Mathieu Simon, Sorina Camarasu Pop, Pascal Girard, Roxana Améli, Jean-Christophe Ferré, Anne Kerbrat, Thomas Tourdias, Frédéric Cervenansky, Tristan Glatard, Jérémy Beaumont, Senan Doyle, Florence Forbes, Jesse Knight, April Khademi, Amirreza Mahbod, Chunliang Wang, Richard McKinley, Franca Wagner, John Muschelli, Elizabeth Sweeney, Eloy Roura, Xavier Lladó, Michel M. Santos, Wellington P. Santos, Abel G. Silva-Filho, Xavier Tomas-Fernandez, Hélène Urien, Isabelle Bloch, Sergi Valverde, Mariano Cabezas, Francisco Javier Vera-Olmos, Norberto Malpica, Charles Guttmann, Sandra Vukusic, Gilles Edan, Michel Dojat, Martin Styner, Simon K. Warfield, François Cotton, Christian Barillot

**Affiliations:** VISAGES: INSERM U1228 - CNRS UMR6074 - Inria - University of Rennes I, France; Department of Radiology, Lyon Sud Hospital, Hospices Civils de Lyon, Lyon, France; LaTIM, INSERM, UMR 1101, University of Brest, IBSAM, Brest, France; Univ Lyon, INSA-Lyon, Université Claude Bernard Lyon 1, UJM-Saint Etienne, CNRS, Inserm, CREATIS UMR 5220, U1206, F-69621, LYON, France; CHU Rennes, Department of Neuroradiology, F-35033 Rennes, France; CHU Rennes, Department of Neurology, F-35033 Rennes, France; CHU de Bordeaux, Service de Neuro-Imagerie, Bordeaux, France; Department of Computer Science and Software Engineering, Concordia University, Montreal, Canada; Pixyl Medical, Grenoble, France; Inria Grenoble Rhône-Alpes, Grenoble, France; Image Analysis in Medicine Lab, School of Engineering, University of Guelph, Canada; Image Analysis in Medicine Lab (IAMLAB), Ryerson University, Canada; School of Technology and Health, KTH Royal Institute of Technology, Stockholm, Sweden; Department of Diagnostic and Interventional Neuroradiology, Inselspital, University of Bern, Switzerland; Johns Hopkins Bloomberg School of Public Health, Baltimore MD; Research institute of Computer Vision and Robotics (VICOROB), University of Girona, Spain; Centro de Informática, Universidade Federal de Pernambuco, Pernambuco, Brazil; Depto. de Eng. Biomédica, Universidade Federal de Pernambuco, Pernambuco, Brazil; Computational Radiology Laboratory, Department of Radiology, Children’s Hospital, 300 Longwood Avenue, Boston, MA, USA; LTCI, Télécom ParisTech, Université Paris-Saclay, Paris, France; Medical Image Analysis Lab, Universidad Rey Juan Carlos, Spain; Center for Neurological Imaging, Department of Radiology, Brigham and Women’s Hospital, Boston, MA, USA; Inserm U1216, University Grenoble Alpes, CHU Grenoble, GIN, Grenoble, France; Department of Computer Science, University of North Carolina, Chapel Hill, NC, USA

**Author notes:** Email address:* (Olivier Commowick).

**Keywords:** Multiple sclerosis, image segmentation, performance evaluation, computing infrastructure, open science, distributed computing

## Abstract

We present a study of multiple sclerosis segmentation algorithms conducted at the international MICCAI 2016 challenge. This challenge was operated using a new open-science computing infrastructure. This allowed for the automatic and independent evaluation of a large range of algorithms in a fair and completely automatic manner. This computing infrastructure was used to evaluate thirteen methods of MS lesions segmentation, exploring a broad range of state-of-the-art algorithms, against a high-quality database of 53 MS cases coming from four centers following a common definition of the acquisition protocol. Each case was annotated manually by an unprecedented number of seven different experts. Results of the challenge highlighted that automatic algorithms, including the recent machine learning methods (random forests, deep learning, …), are still trailing human expertise on both detection and delineation criteria. In addition, we demonstrate that computing a statistically robust consensus of the algorithms performs closer to human expertise on one score (segmentation) although still trailing on detection scores.

## 1. Introduction

Multiple Sclerosis (MS) is a chronic inflammatory disease of the central nervous system affecting around 2.5 million persons worldwide, with a prevalence rate of 83 per 100000 (higher rates in countries of the northern hemisphere) and a woman:man ratio of around 2.0 [1]. It is characterized by widespread inflammation, focal demyelination, and a variable degree of axonal loss. With the appearance of new treatment molecules modifying the disease evolution (disease modifying drugs - DMD), one of the major challenges in treating multiple sclerosis is now to overcome classical clinical criteria, such as the expanded disability status scale (EDSS), to go towards more sensitive and specific criteria. In this context, Magnetic Resonance Imaging (MRI) plays an important role for the diagnosis [2] and evaluation of the evolution of the disease, thus providing insights to adapt the treatment to each individual due to the highly variable nature of the MS disease course [3].

In this context, the number and spread of lesions in the patient’s parenchyma (and their evolution) [2] has become a crucial information on the patient’s disease status, which may then be used for validating the patient treatment. This task however requires the delineation of MS lesions: a tedious, manual operation performed by the radiologist. In addition, this delineation is prone to inter-expert variability, especially when the images being used for segmentation differ from a center to another (in terms of protocols, modalities and intrinsic MRI quality). Doing this task manually on large databases of patients is therefore almost impossible and automatic algorithms, thoroughly validated, have become a crucial need for the clinical community. To simplify the clinician’s task, a large literature of automatic segmentation methods has been devised [4, 5, 6] with a large spectrum of algorithms from classical tissue intensity classification and lesion modeling to machine learning.

All published approaches are however evaluated on different datasets, usually not calibrated, and their results are therefore usually not directly comparable, making difficult the choice of the most relevant method adapted to a clinical context. To overcome this issue, competitions (so-called challenges) have been organized in MS lesion segmentation in the past years. The first one was organized at the MICCAI 2008 conference [7]. It evaluated nine different methods on a database of 45 patient images (from two different centers: 20 for training and 25 for testing), with respect to a ground truth composed of two expert segmentations for each case. However, no protocol standardization was performed between the two sites, therefore two raters were not enough to handle the variability in the acquired images and get a sufficiently reliable consensus manual segmentation. The second major challenge on MS lesion segmentation was held in 2015 at the IEEE ISBI international conference [8]. It was more focused on the study of longitudinal lesion evolution with specific evaluation metrics based on segmentation volume evolution (in addition to the regular segmentation overlap metrics used in 2008). This challenge evaluated 10 different methods on a dataset composed of five patients images each with an average of 4.4 time points, each time point being manually delineated by two experts. As for the MICCAI 2008 challenge, two raters were not enough to account for disparities in the different raters manual segmentations and get a representative consensus. The process of evaluation was similar for the two challenges. A subset of the patient images was provided to the participants with the ground truth (GT) segmentation to the participants for them to train their respective methods. In a second step, a testing set was provided (without the ground truth) to the participants asking them to submit back their results. Evaluation was then performed on those results using overlap-based metrics.

Several problems may however affect such challenges. First, as a general comment for all challenges, a lack of fairness may exist between the participants: since the testing images are provided, some participants may indeed optimize the parameters of their algorithms on a patient basis to obtain better results. Doing this illustrates the potential of the method but not its practical usability: a clinician would prefer to use always the same set of parameters to process each new or returning patient. In addition, since participants run their algorithm on their own computing environment, no evaluation relative to computing performance (e.g. required memory of computing time) is possible. There is therefore a need for computing platforms for supporting challenges including data storage, processing pipelines (i.e. segmentation algorithms work-flow used) integration and evaluation on stored datasets. Such platforms would provide a truly fair comparison between fully automatic methods. In addition, such remote computing platforms, able to host a large variety of algorithms, announce what the future cloud computing services will provide to assist clinicians (radiologists, neurologists, …) in using computer aided diagnosis solutions. This computing environment also opens the road to open-science platforms where people will find solutions to post their data, send or retrieve algorithmic solutions and provide an independent yet secure environment to compare, assess and combine various algorithms outcomes and solve clinical problems.

Another issue in segmentation challenges is the number of manual delineations to compute the ground truth. Usually only two are available, which is insufficient to illustrate the inter-expert variability, particularly when considering MS lesions segmentation. Finally, and specifically to MS lesions segmentation, previous challenges considered only segmentation based metrics, ignoring the number of correctly detected lesions independently of their shape, which is an acute criterion to assess the disease evolution [2]. This would be very beneficial for the clinician, especially when considering MS evolution where the number of new lesions is critical.

We proposed and organized in 2016 a new generation of segmentation challenge hosted at the MICCAI international conference **(http://www.miccai2016.org)**. It aimed at proposing solutions to several of the previously mentioned defects first by gathering an unprecedented database of MS patients, coming from three different centers (representing four different scanners, one of which was intentionally hidden at the training phase from the challengers to test their algorithms’ adaptation capabilities) but all following a common consensus protocol [9], each patient being delineated by seven experts to evaluate not only automatic methods performance but also inter-expert variability of manual segmentation. We have performed the evaluation on a dedicated computing platform provided by France Life Imaging (**https://www.francelifeimaging.fr/en**), providing pipeline integration, database storage and automatic execution capabilities. Challenge participants were asked to train their algorithms on a reduced set (*n* = 15) and then integrate their pipeline on the platform, requiring no action from them in the latter parts of the evaluation process (*n* = 38). We also proposed an evaluation strategy on two separate levels: a segmentation level where the overlap precision of the segmentation was evaluated; and a detection level where the number of correctly detected lesions was evaluated, independently of the precision of their shape.

We present in this article a retrospective analysis of this challenge and the methods we used to obtain those results. The main outcomes of the challenge highlighted that automatic algorithms are still trailing human expertise on the front of MS lesions segmentation and sensitive to unknown images (different scanners) even with an harmonized acquisition protocol. This happens for all methods, independently of their category (recent machine learning algorithms including deep learning or random forests or more classical tissue classification algorithms). In addition, we demonstrate how using an open-science computing environment allows for the combination of multiple algorithmic outcomes, and how combining these algorithms could lead to improvements in detection and contouring of MS lesions. Together with the computing platform introduced in this paper, this could lead to tremendous help for the clinicians in the use of automatic segmentation algorithms to support their diagnosis and treatment follow-up in MS.

## 2. Results

### 2.1 Challenge data, computing platform and participating teams

The first major result of this study is the gathering of a database of 53 multiple sclerosis patients with “ground truth” of very high-quality. The database patient scans were following the OFSEP protocol recommendations in [9], which is currently applied in France for the constitution of the national cohort in MS (for more details on the protocol, see Section 4.1). Following this approach has allowed for an evaluation representative of the current imaging protocols standards and easily usable to characterize the best performing algorithm for future use. This standardization of imaging protocols announces how the dissemination of computer aided diagnosis and imaging biomarkers solutions will be implemented in the future. Image processing algorithms indeed need image normalization and quality control to ensure peak performance. In addition, the images came from three different sites in France on four different MRI scanners and different manufacturers (Siemens, Philips and GE) including three 3T and one 1.5T magnets. For each MS patient case, an unprecedented number of seven manual delineations was gathered, from trained experts split over the three sites providing MR images. From these segmentations, a consensus “ground truth” segmentation was built for evaluation with the LOP STAPLE algorithm [10]. We present in Fig. 1 an example of a patient 3D FLAIR, the seven manual segmentations of lesions and their consensus segmentation, illustrating the variability for a representative patient between expert segmentations. Patients demographic data were the following: average age of 45.3 years (± 10.3 years) with a male:female ratio of 0.4. This database was then split into two sets: one training set of 15 patients from three scanners (thus intentionally missing one scanner from the database) given to participants, and one testing set of 38 patients, not seen by the participants, used for evaluation. Demographics of patients do not vary significantly over the different sitesin terms of age. Some variations exist in the male:female ratios in some centers. The training and testing sets have an average age difference of 5 years (training set patients are 5 years younger).

**Figure 1:**
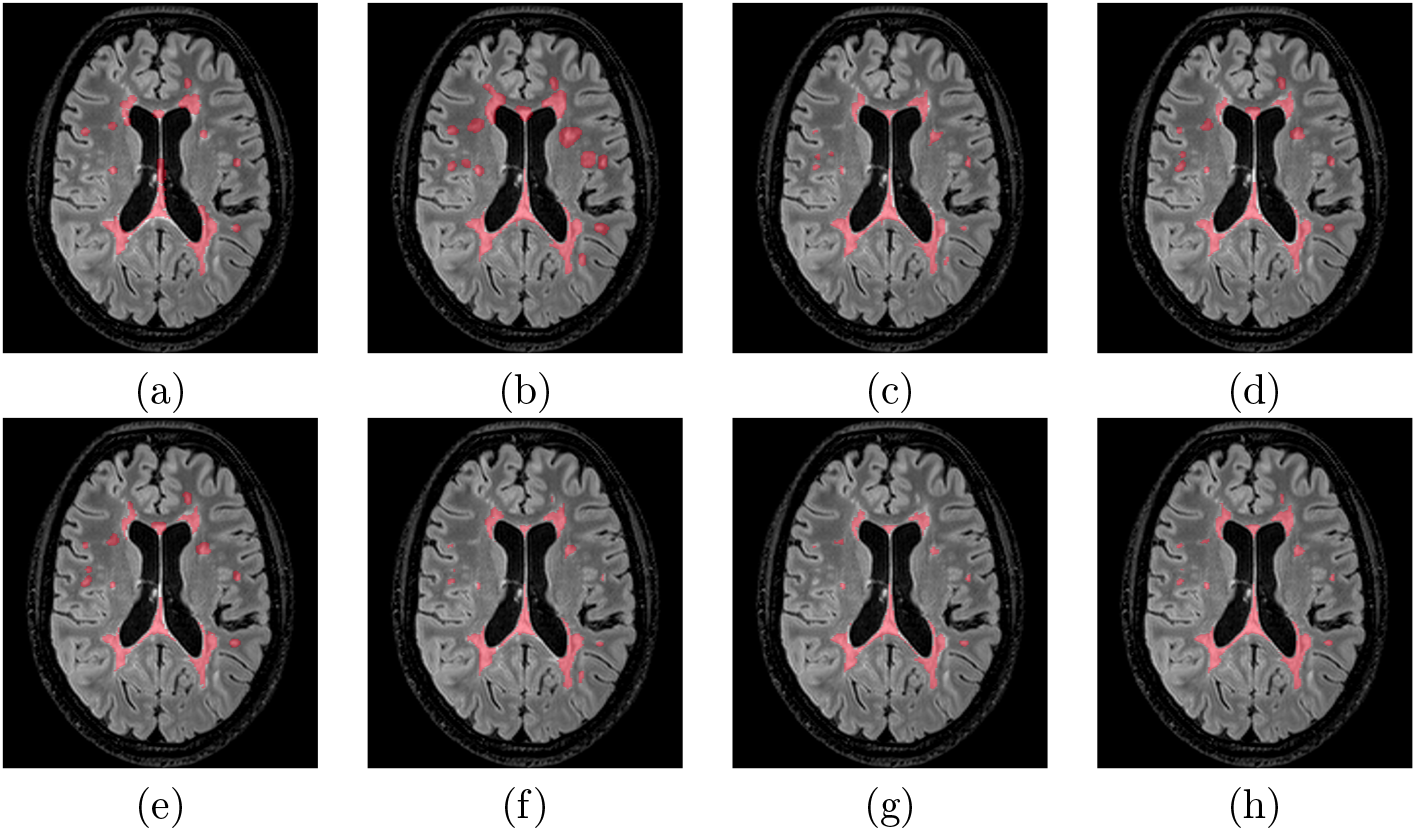
Illustration of an MS patient delineations overlaid on the 3D FLAIR image. (ag): individual manual delineations of MS lesions from each of the experts, (h): consensus segmentation considered as the ground truth.

A total of thirteen teams were evaluated, and the website (http://portal.fli-iam.irisa.fr/msseg-challenge/), databases and algorithms will remain open for future use. A summary of the evaluated methods is presented in Table 1 with a short description of their characteristics (MR sequences used as input, implementation, main methodology). The algorithms evaluated in the challenge are representative of a broad range of the available methods in the recent literature, with unsupervised tissue classification methods, level-sets, random forests and deep learning (convolutional neural network, artificial neural networks). Depending on the challenger team, the image modalities used for the segmentation varied from just one (usually FLAIR) to all provided modalities. Most evaluated algorithms ran on regular computer CPU, while two (team 6 and 12) leveraged specific hardware (GPUs) for intensive computation (e.g. deep learning). The computing infrastructure was able to provide the relevant computing solution for all requirements.

**Table 1:**
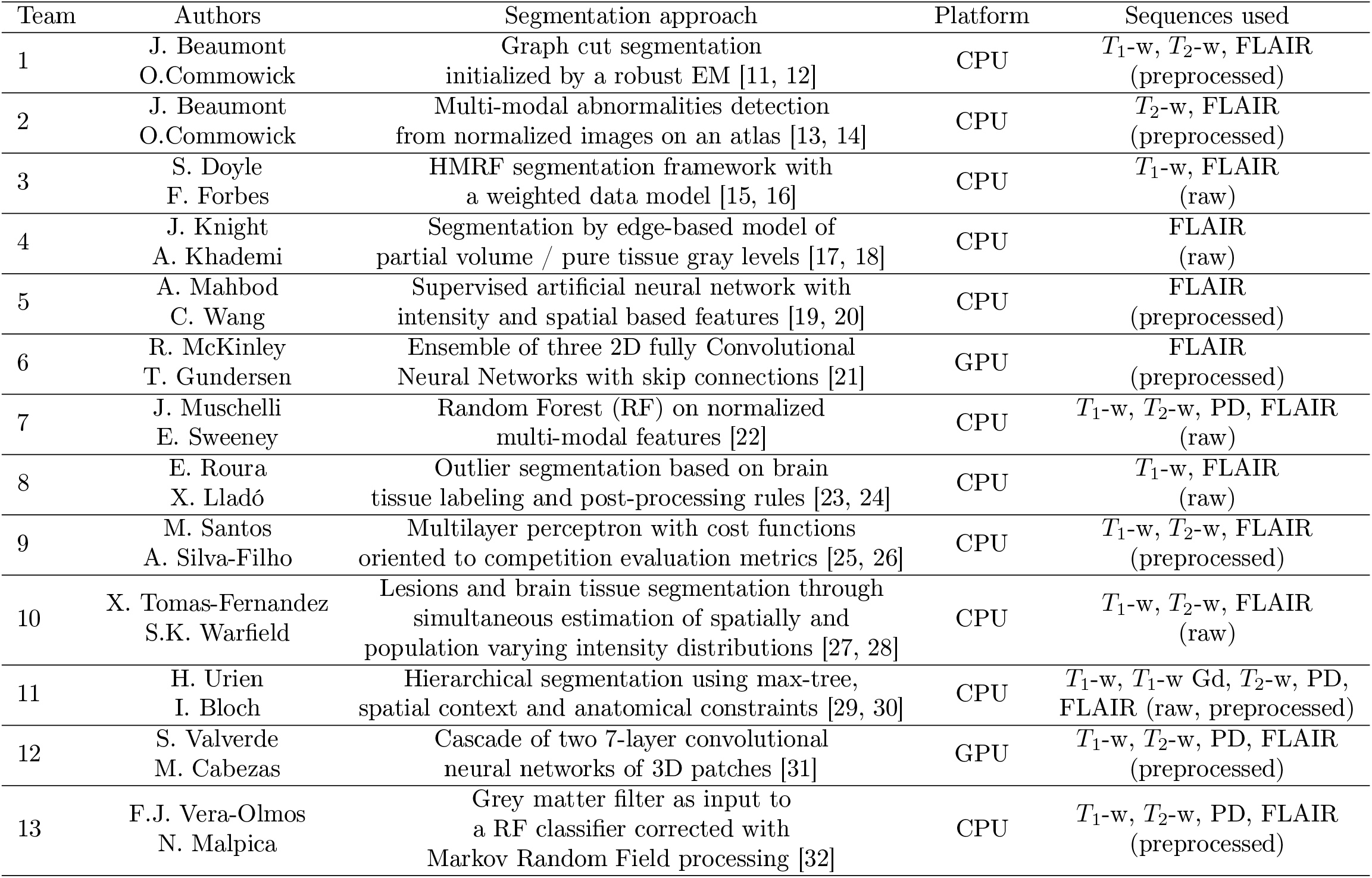
MS lesion segmentation methods evaluated at the MICCAI 2016 challenge.

These teams were evaluated in a distributed Web platform based on software containers provided by the France Life Imaging platform, allowing for the automatic, challenger independent evaluation of the algorithms (see Section 4.3 for more details on the challenge execution platform). Using such a platform, providing integrated storage and computing facilities for challenges, allowed for fair comparisons as challengers could not tune their algorithm specifically for each test patient. Each challenger was indeed asked only to provide a binary image (i.e. an annotated Docker container image) of their processing pipeline and the evaluation was later on run automatically on the platform with the following metrics used.

### 2.2 Two kinds of performance metrics were set up for evaluation

Clinicians evaluate lesion segmentation in multiple sclerosis with different criteria. Lesion segmentation precision, i.e. the precision of contours delineated for each lesion, is crucial as the total volume of lesions (total lesion load - TLL) is part of the criteria to evaluate disease severity [33, 34]. When coming to pathology evolution or treatment efficiency evaluation however, lesion count and particularly the number of new lesions independently of their sizes is key. Moreover, this lesion count is a crucial component of MS diagnosis according to McDonald criteria [2]. For these tasks, detecting all lesions is more important than their precise contours. We have therefore implemented a large set of evaluation measures for the challenge with the goal of evaluating these different aspects. Evaluation in the following is therefore split into three major categories of evaluation metrics:

- Segmentation evaluation: does the algorithm provide a precise delineation of each lesion? This category includes average surface distance and Dice overlaps as the main metrics
- Lesion detection evaluation: does the algorithm find all lesions in the image independently of its precise delineation? This category includes the *F*_1_ score, gathering in one scalar information on the number of lesions correctly and incorrectly detected

### 2.3. All methods are outperformed by the experts

We have automatically clustered the average algorithms and experts annotations agreements (with their covariances accounted for) with respect to the “ground truth” (see Section 4.4 for more details). Results of this clustering, illustrated in Fig. 2 for all couples of measures considered in the challenge, highlight a major result of the challenge: over all patients and all evaluation metrics, each individual method performs slightly below all experts. On all graphs in Fig. 2, all experts (and only them) are indeed always grouped in a single cluster that performs better than all automatic algorithms. Two other clusters are also distinguished in these graphs, which vary depending on the evaluation metric, that regroup better performing and lower performing algorithms for each couple of evaluation metrics.

**Figure 2:**
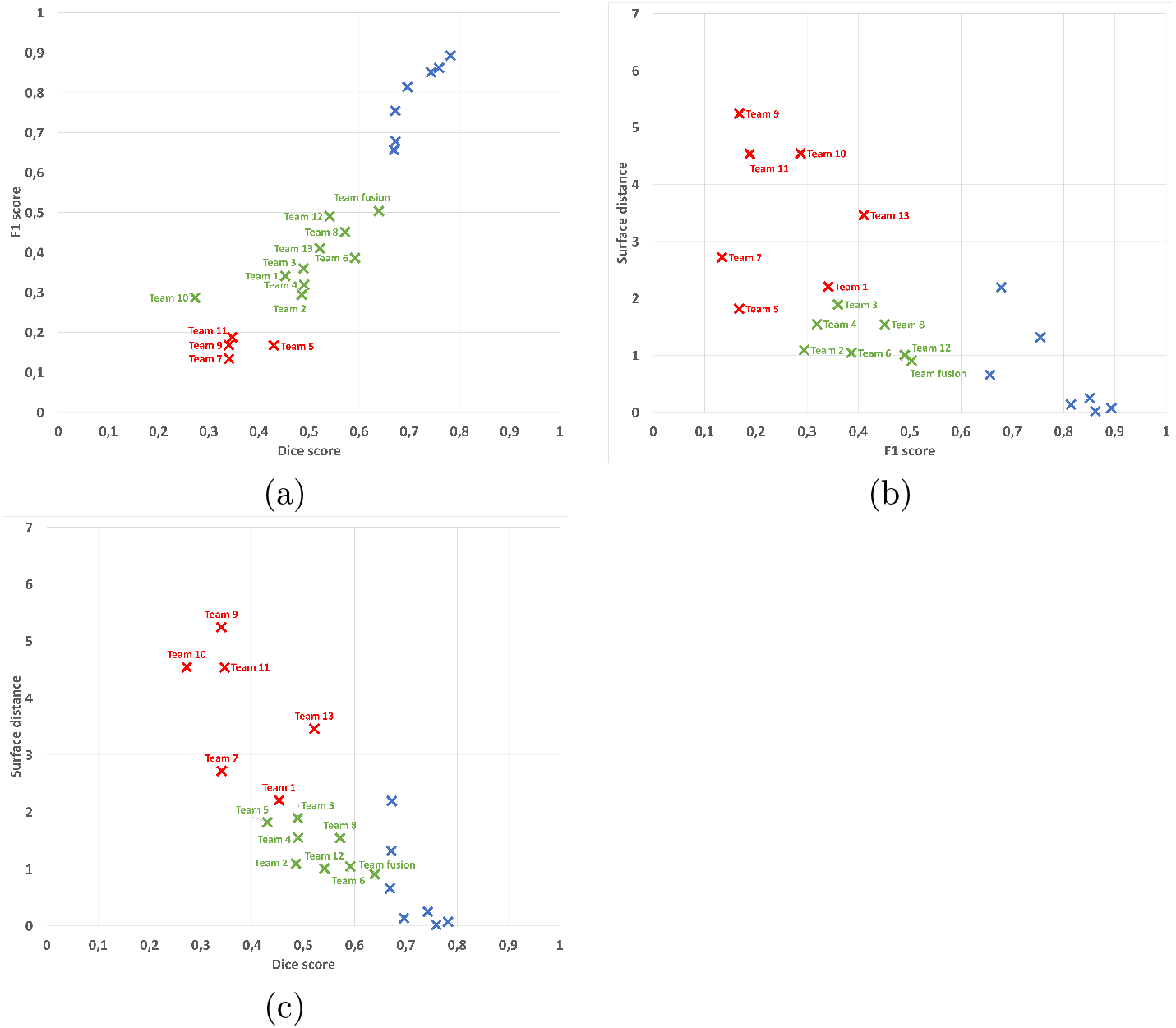
Graphical results illustration of automatic clustering of average results for each team and expert into three groups (scatter plots of pairs of two evaluation parameters: (a): Dice and *F*_1_ scores, (b): surface distance and *F*_1_ scores, (c): surface distance and Dice scores). Legend: blue crosses: group 1 (always containing only the seven experts even though the clustering is automatic), green crosses: group 2 (best performing algorithms), red crosses: group 3 (lower “quality” algorithms). Team numbers associated with each point on the graph are indicated as labels. Team fusion indicates a composite segmentation result further discussed in Section 2.5.

In those graphs, we can additionally study the performance of automatic algorithms with regards to each evaluation metric considered (average surface distance, Dice score and *F*_1_ score). Automatic methods fail much more on the detection of lesions (F_1_ score), with a minimum average score of 0.13 and maximum average of 0.49, while the minimum average score obtained by an expert is 0.66 (significant difference, Wilcoxon signed rank test, *p* = 3.7 × 10^−5^). This is understandable however as all algorithms are primarily designed to obtain the best segmentation scores while not considering lesion detection which is a somewhat different task. However, even on the Dice score, which is a segmentation metric, the best automatic method performs lower than the lowest expert average score: it reaches an average of 0.59 while the lowest expert is on average at 0.67 (significant difference, Wilcoxon signed rank test, *p* = 2.9 × 10^−3^). The average surface distance is a more balanced metric in terms of results with the second group of algorithms in each graph reaching the level of agreement that the experts do with the consensus.

### 2.4. Segmentation on an unknown scanner leads to poorer performance

Scanner 3 in the testing database was unknown to the teams participating to the challenge. On this center, we have evaluated how automatic algorithms performed without knowing the image characteristics beforehand. The results of this comparison highlight for a large number of automatic algorithms a slight decrease in performance when encountering unknown images, even if they come from a common protocol. This evaluation per center and per evaluation metric is presented in Fig. 3.

Looking closer at the graphs in Fig. 3, we can observe for the detection metric (*F_1_* score, Fig. 3.b) a slight decrease for 8 teams among the thirteen evaluated, leading to an average score over all teams of 0.22 for center 3 while the same automatic methods range between 0.32 and 0.39 for other centers. The same trend can be observed for the Dice score segmentation metric (Fig. 3.a) with a slight decrease in performance for 8 teams as well, and an average score over all teams of 0.38 while the same algorithms reach a level ranging between 0.48 and 0.50 on other centers. The observations are different for the average surface distance (Fig. 3.c), where results for center 3 are at the same level as center 8, however the variance in results is much higher, preventing from finding any statistically significant difference on that metric.

### 2.5. Combining methods through label fusion improves over individual algorithms

In addition to the individual automatic algorithms, we have evaluated a composite team named “team fusion”. This method gathered the other thirteen teams segmentations in a consensus through label fusion using the LOP STAPLE algorithm [10]. The goal of this fourteenth method was to evaluate the capability of such a label fusion method to overpass the individual difficulties of each method and thus obtain results closer to the ground truth. We present the results of this evaluation on the different evaluation metrics in Fig. 4.

This composite algorithm improves the average results the average results of individual automatic algorithms for all metrics, suggesting its ability to incorporate the best of each team into a consensus segmentation, better in line with the experts. These results are confirmed by points “Team fusion” in the clustering graphs in Fig. 2. However, the results obtained are still not perfect and lag behind the experts level of agreement with the “ground truth”. More precisely, the improvement of team fusion over other algorithms is particularly visible on segmentation metrics (Dice scores and average surface distance) since it provides segmentation performances similar to the lowest experts. This improvement is however less important on the detection metric (*F_1_* score). This smaller improvement seems logical as the label fusion algorithm used for team fusion is primarily designed to optimize segmentation performance and not specifically detection. With that said, the first position of Team fusion among the segmentation methods illustrates how a composite algorithm mixing results of other teams is able to perform better than each individual automatic method. This also illustrates the importance to provide an open-science computing platform able to combine results of independent algorithms.

**Figure 3:**
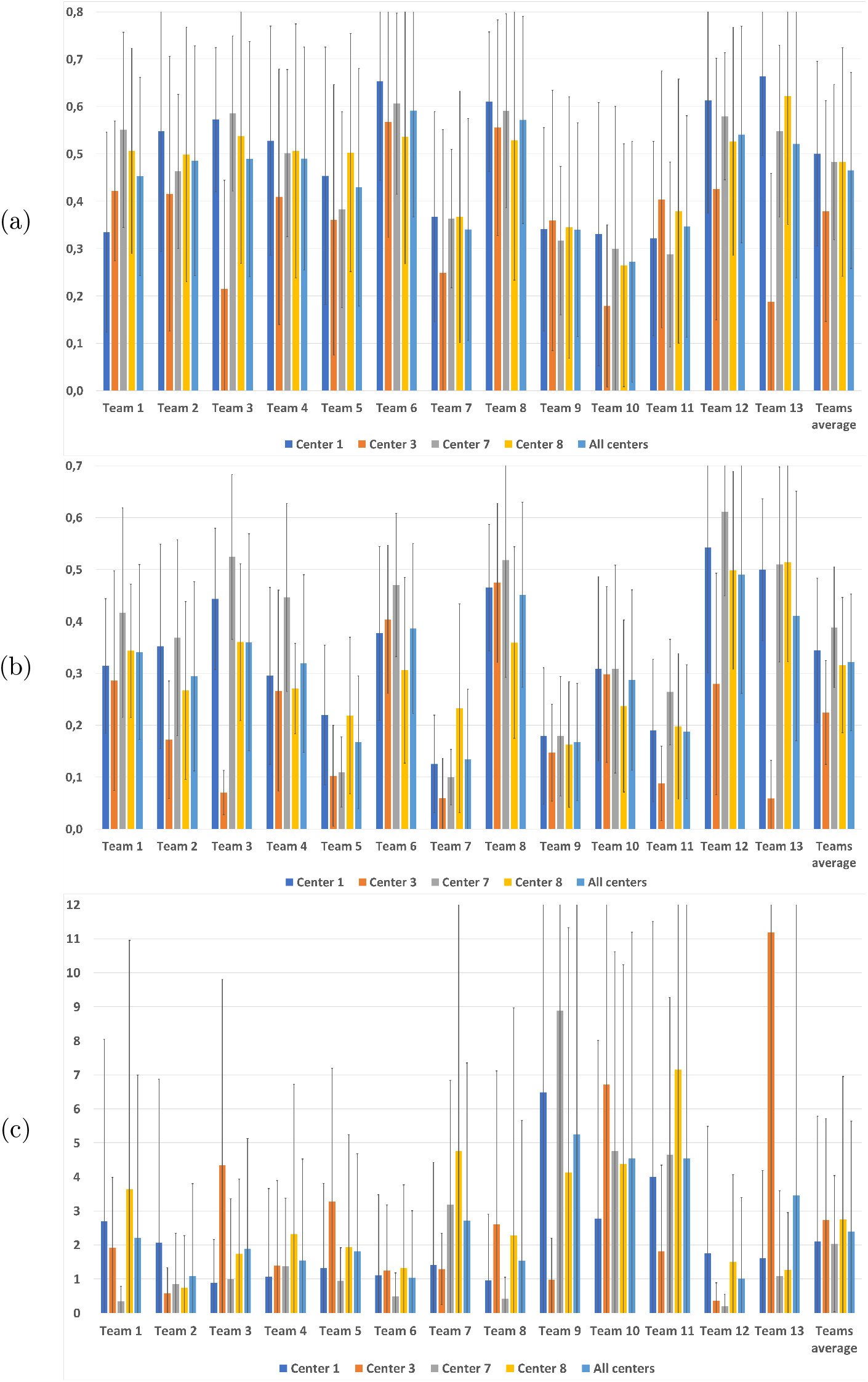
Dice scores (a), *F*_1_ scores (b) and average surface distances (c) with respect to the consensus per team for each center and averaged over all centers.

**Figure 4:**
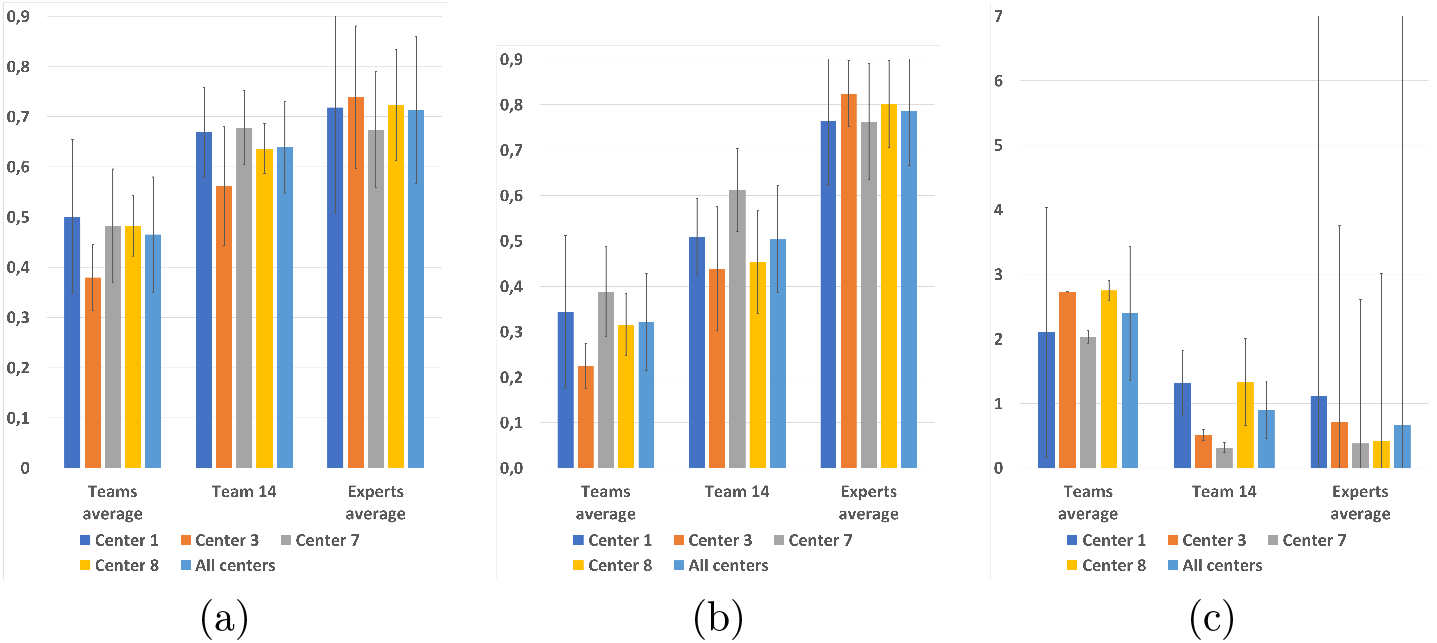
Dice scores (a), *F*_1_ scores (b) and average surface distances (c) with respect to the consensus for each center and averaged over all centers for composite Team fusion with respect to the average experts agreement level.

### 2.6. Lesion load and lesion size directly influences automatic segmentation quality

We additionally performed an experiment to evaluate, independently of their individual behaviors, the algorithms sensitivity to the true amount of lesions in the “ground truth” for a given patient. To this end, we averaged the Dice scores (respectively the average surface distances and *F*_1_ scores) over all methods for each patient and plotted in Fig. 5 this average value with respect to either the number of lesions or the total lesion load in the consensus. On each graph, we then computed a log-linear regression for which we display the Spearman squared correlation.

From Fig. 5, it is clear that the worst results are obtained for patients whose total lesion load is low. This is especially true for segmentation performance scores: the Dice score (the squared correlation *R^2^* of the regression reaches 0.82) and the average surface distance (*R^2^* of 0.71). For the *F*_1_ score (a detection metric), the correlation is however weaker than for the total lesion load (*R^2^* of 0.45). From these graphs, the correlation between the number of lesions and the obtained scores is less clear, all correlations being smaller than with the total lesion load (*R^2^* of 0.46 with the Dice score, 0.38 with the *F*_1_ score, and 0.60 with the average surface distance). This result however seems reasonable since a patient presenting many small lesions is intuitively more difficult to delineate than a patient with a small number of large lesions.

**Figure 5:**
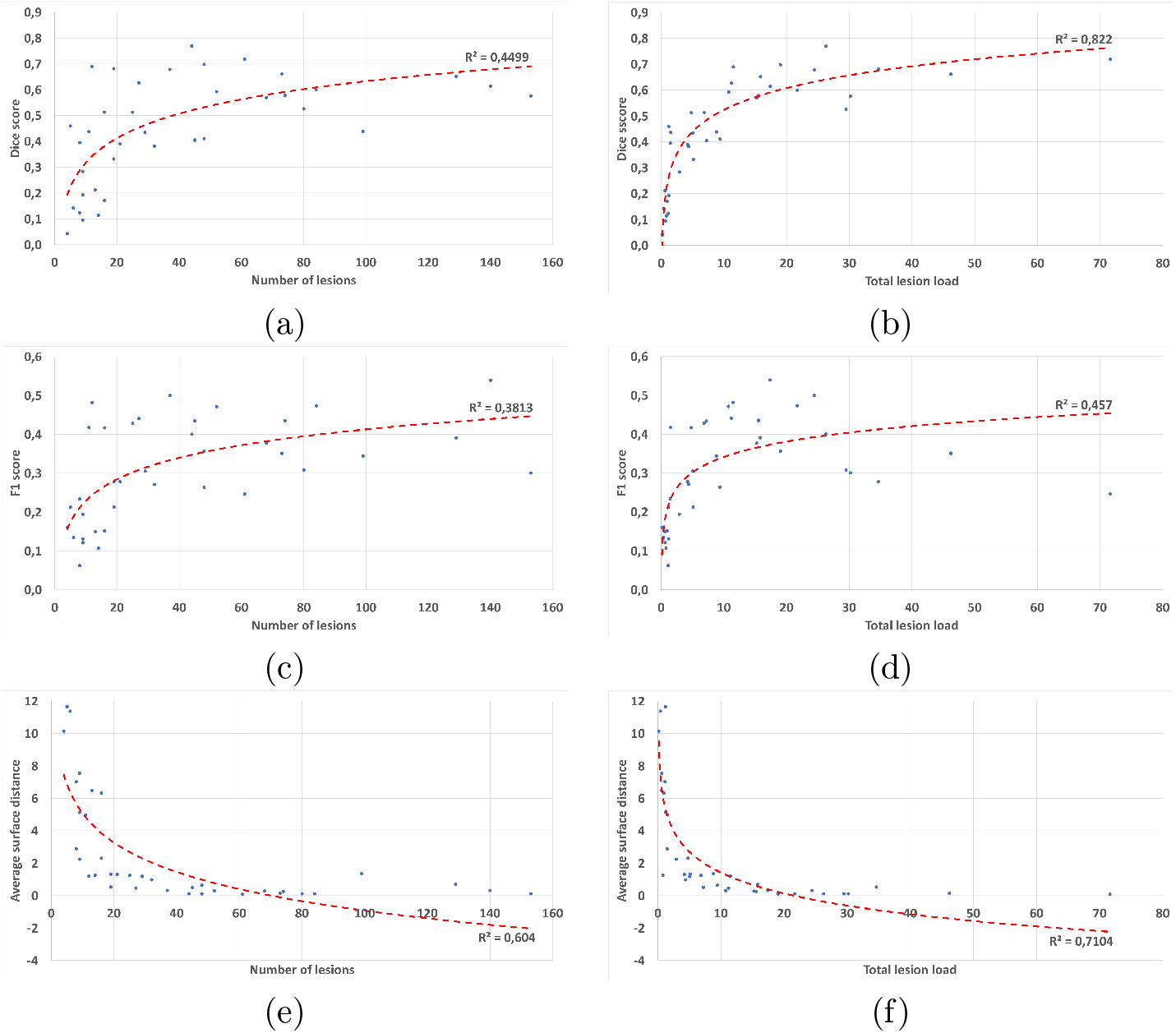
Link between average scores of all methods and number of lesions (first column) and total lesion load (cm^3^, second column). First line: Dice score, second line: *F*_1_, third line: average surface distance.

Total lesion load in a patient is thus very correlated with segmentation and detection scores while not with the number of lesions. To further qualify this fact, we performed an experiment considering detection scores individually for each lesion in regard of its volume. We have thus computed, for each team and for each lesion of the “ground truth” of each patient, a binary detection score telling whether the lesion was detected or not by a specific team. Counting the number of teams which detected the lesion thus provides us with a rate of detection for each lesion (a rate of 0% meaning that no team detected the lesion, and 100% meaning that all teams detected the lesion). Those detection rates were further binned according to lesion volume. The graph in Fig. 6 illustrates their relation with respect to lesion volume (in mm^3^).

**Figure 6:**
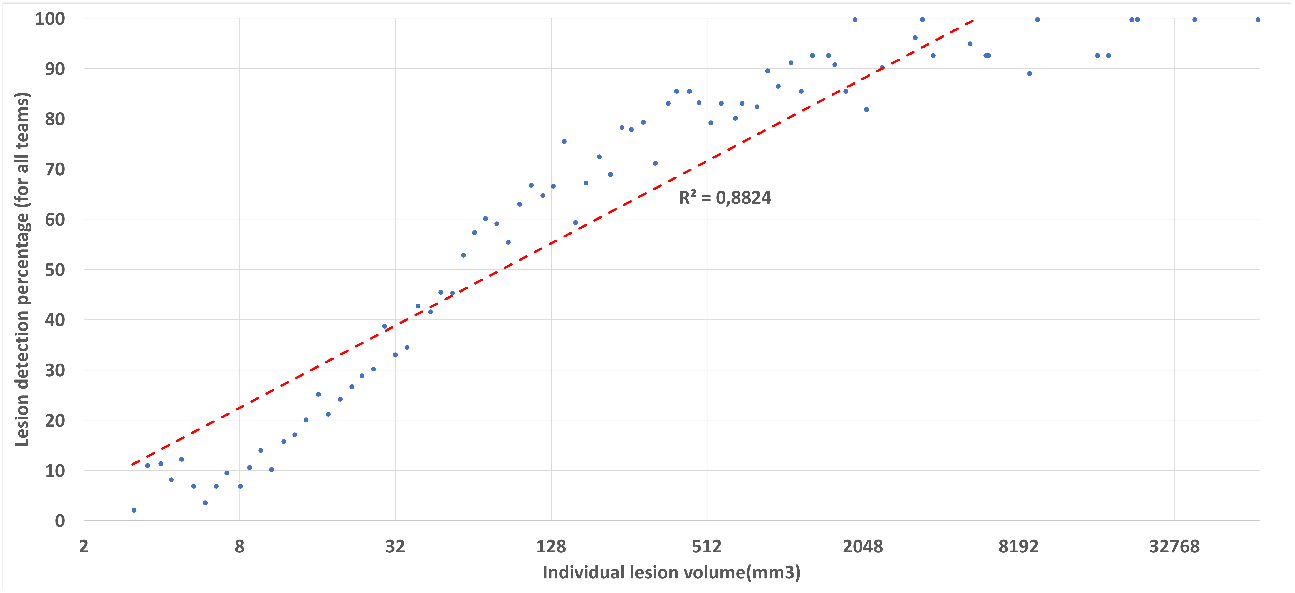
Individual lesion detection rate (average over all methods) as a function of lesion size. X-axis: individual lesion volume on a logarithmic scale. Y-axis: detection rate (in percentages, number of teams detecting a lesion of this volume over all patients).

From Fig. 6, we can clearly see that not only total lesion load influences segmentation quality, but lesion volume is also clearly linked with lesion detection (*R^2^* of 0.88 after a logarithmic linear regression). All methods tend to fail (rates of detection going to zero) for small lesions, while almost all teams are detecting the lesions when their volume is sufficiently large.

### 2.7. Delineating an image with empty consensus

We finally present in Table 2 results obtained by each expert and each evaluated pipeline on a specific case, from Center 7, where no lesion was present in the consensus segmentation. In this specific case, none of the proposed detection and segmentation measures can be used since they all rely on the fact that the consensus is not empty. We thus instead defined two specific metrics: the number of lesions detected (i.e. the number of connected components whose size is larger than 3 mm^3^ in the segmentation), and the total lesion load (i.e. the total volume of the previously extracted connected components) found by each algorithm. For both metrics, since the consensus contains no lesions, a perfect value is 0 while the results get worse when the metrics grow.

**Table 2:** Number of lesions and lesion volume detected by each team and expert on the no consensus lesion case.

| | Lesions volume ($\text{cm}^3$) | Number of lesions |
| --- | --- | --- |
| Expert 1 | 0.052 | 2 |
| Expert 2 | 0.090 | 2 |
| Expert 3 | 10.887 | 8 |
| **Expert 4** | **0** | **0** |
| Expert 5 | 0.017 | 1 |
| **Expert 6** | **0** | **0** |
| Expert 7 | 0.029 | 2 |
| Team 1 | 8.252 | 18 |
| **Team 2** | **0** | **0** |
| **Team 3** | **0** | **0** |
| Team 4 | NA | NA |
| Team 5 | 28.436 | 522 |
| Team 6 | 0.473 | 7 |
| Team 7 | 5.990 | 168 |
| **Team 8** | **0** | **0** |
| Team 9 | 2.545 | 33 |
| Team 10 | 11.085 | 31 |
| Team 11 | 3.436 | 42 |
| Team 12 | 0.056 | 1 |
| Team 13 | 0.074 | 4 |

Several observations may be drawn from this table. First of all, among experts, two delineated no lesions while five actually delineated one or several lesions (from one to 8 depending on the expert, and from 0.02 to 11 cm^3^). The fact that the consensus is empty therefore means that the experts were not agreeing on the position and extent of lesions, which lead to no lesions in the final “ground truth”. Among the automatic segmentation pipelines, the results are also largely varying: depending on the team, the number of lesions delineated varies from 0 to 522, while the lesion load detected varies from 0 to 28.44 cm^3^. In addition, this image caused problems to some algorithms not initially designed for patients without lesions (team 4). This is an interesting case as it highlights the different behaviors of the algorithms on a case for which the pipelines were not designed. Overall, we can notice that most of the methods behave well in comparison to the experts.

## 3. Discussion

We have presented the first challenge based on an integrated computing platform, applied to multiple sclerosis lesions segmentation. The challenge computing platform was constituted of 1- a database to store the challenge images and results from the challengers, 2- a computing platform on which the evaluation was performed independently of the participants who were asked to post their processing pipelines, and 3- an automatic evaluation of the results against the “ground truth”. This open-science computing platform has many advantages, including a fair comparison of the participants algorithms being run on the same platform and with the same set of parameters for all patients. In addition, the packaged algorithms may be re-used for other applications or if the validation database gets extended. As future work, we plan at transitioning to use the BIDS format (http://bids.neuroimaging.io) to provide a standardized and more intuitive way of storing both the input data and output results. This would provide a great improvement in easing the pipeline design and integration in the platform.

This platform was put together for the specific organization of a challenge on multiple sclerosis segmentation with a database of 53 patients each with an unprecedented number of seven manual segmentations from trained experts. A total of thirteen teams participated, illustrating the variety of algorithms both in terms of methodology and implementation. All results computed from the challenge were very insightful and revealed several points worth of discussion. First of all, despite that methods vary in their results, the point where a single automatic method is able to perform as well as the consensus of the experts has not yet been reached. The experts are indeed always slightly better than any method for all performance measures. More specifically, all methods perform relatively poorly on detection metrics, which is however an important point for MS diagnostic and clinical evaluation of the patient evolution. Historically all methods have been interested in segmenting well the contours of the lesions rather than counting well the number of lesions. As a consequence the detection metrics are not optimal, which explains why results are well below the experts. On a more positive point, recent methods such as those based on machine learning (especially deep learning) have made great progress and the gap is reducing, which leaves hope to reach the same level of agreement than the experts. In addition, it should be remembered that it is always difficult to define a “ground truth” for MS lesions segmentation. The experts have indeed a relatively large variability, which comes from different appreciations of the image and of the definition of a lesion. Finally on this point, it is also interesting to note that a composite method for segmentation (team fusion) combining the different automatic methods while rejecting outliers is able to drastically improve segmentation results and get closer to the ground truth. However, this happens mainly for segmentation performance metrics and less for detection performance metrics. This may be due to the inherent design of the label fusion methods that do not work on a lesion basis but rather on a voxel basis, and thus favor segmentation based metrics. In addition, since many methods fail at delineating well lesions when the total lesion load is small (see Fig. 5), this composite method is not able to perform a good segmentation for these cases.

We have also illustrated through this challenge the behavior of methods in several specific cases: testing for scanner dependency of the algorithms, where we compared the results for four different scanners, one of them being hidden from the training dataset. This study illustrated that all methods are still sensitive to scanners on which they were not trained for (either training in the machine learning sense or training in the sense of parameters tuning by a human being), obtaining lower scores for those images. On the contrary, when the training set included representative images of different scanners, all algorithms behaved equally independently of the center or scanner. This suggests the importance of looking for more training independent methods or, meanwhile, to have enough representative cases to train on (at the high cost of providing multiple manual image annotations from experts). A second specific case considered a patient with no lesions in the ground truth provided from the experts. For that patient, even if some methods had not planned this case, all methods behaved globally well even though participants were not told in advance about this fact.

While not mentioned explicitly in this article, we also looked at the relation between algorithm performance and preprocessing or modalities used. For both aspects, there is no clear evidence of a link. Some algorithms perform well while using only a subset of modalities, while some other that use all modalities perform less well. However the reverse is also true: some of the best algorithms use all modalities while some less good ones use a smaller number of modalities. This is however a crucial aspect as information on this could help in the design of shorter acquisition protocols in the future, using only those modalities useful for automatic segmentation. Future works, which have to be in close link with the challenge participants (but facilitated thanks to our computing platform), will look at the robustness of results of the individual algorithms with respect to both preprocessing used and modalities used. This will provide a great insight into optimal, fast protocol design and optimal preprocessing.

We have clearly demonstrated in Fig. 5 a link between the total lesion load in a patient and the performance of segmentation methods: the smaller the true total lesion load, the worse the segmentation results were for every metric. As mentioned earlier, this is partly linked to the fact that voxel-based performance metrics are much more sensitive when the number of voxels in the true segmentation is small. However, this is not the only reason: automatic segmentation methods are indeed behaving slightly worse on these cases and a focus on them should probably help in designing algorithms adapted to all situations. This link between performance and lesion load does not generalize to the number of lesions (also seen in the same figure), which illustrates that there is no clear sensitivity of the methods to the number of lesions for a patient. However, this link is clearly related to a correlation between lesion volume and lesion detection rate, as demonstrated in Fig. 6. This further indicates that lesions are clearly less well detected or even not at all when they are small, which seems rather logical as it intuitively seems tougher for an algorithm to properly locate a small lesion than a larger one.

Evaluation metrics presented in this article are a selection per category (segmentation and detection) of the metrics described in Section 4.4 and computed for the challenge. We chose them as being representative and most informative of the main qualities and defaults of the algorithms. Of course, as illustrated for more general segmentation evaluation purpose [35], three metrics may not be enough for describing the behavior of each method in its entirety. As explained in Section 4, we have complemented these three measures with many other complementary measures, for which we encourage the interested reader to look at the supplementary materials (**http://dx.doi.org/10.5281/zenodo.1307653**).

## 4. Methods

We present in the following the methodological details that allowed us to draw the results and conclusions previously outlined. This section is split in several subparts. Sections 4.1 and 4.2 present in more details the evaluation database used in the challenge and the way in which the manual delineations were carried out and averaged into a “ground truth”. Section 4.3 then outlines the computing platform used in the challenge which was necessary to guarantee a fair comparison of the algorithms. Finally, Section 4.4 presents the evaluation metrics used in the challenge as well as the analyses plan derived from them in this article.

### 4.1. Reference Images Database

In this segmentation evaluation challenge, we relied on a database of images of 53 multiple sclerosis patients following the OFSEP protocol recommendations in [9], which is currently applied in France for the constitution of the national cohort of MS patients. Following this approach allows for an evaluation representative of the current standards and easily usable for newly acquired images. For our challenge, images came from three different sites in France on four different MRI scanners from different manufacturers (Siemens, Philips and GE) including three 3T and on 1.5T magnets. The repartition of the 53 patients is shown in Table 3. More demographic details on the ages and gender repartitions into the training and testing groups are provided in supplementary material (**http://dx.doi.org/10.5281/zenodo.1307653**). Overall, no significant difference of age can be seen between the different centers. While gender differences exist between some centers (in particular center 8), we believe this is of little importance with regard to lesion segmentation and detection quality compared to scanner to scanner differences.

**Table 3:** Demographics data of multiple sclerosis patients collected for the challenge and their repartition among training and testing datasets.

| Center number | Scanner model and site | Training cases | Testing cases | Age (y.o.) | Gender ratio M:F |
| --- | --- | --- | --- | --- | --- |
| 1 | Siemens Verio 3T<br>(University Hospital of Rennes) | 5 | 10 | 43.6<br>± 12.6 | 0.36 |
| 3 | General Electrics Discovery 3T<br>(University Hospital of Bordeaux) | 0 | 8 | 48.9<br>± 11.5 | 0.14 |
| 7 | Siemens Aera 1.5T<br>(University Hospital of Lyon) | 5 | 10 | 45.3<br>± 9.8 | 0.25 |
| 8 | Philips Ingenia 3T<br>(University Hospital of Lyon) | 5 | 10 | 45.5<br>± 7.8 | 0.87 |

These patients were selected to have variable amounts of lesions both in volume and number, and from different centers to represent the variability that may be encountered across sites. For each patient, the following images were provided for each MS patient (see details of the sequence parameters in Table 4): a 3D FLAIR sequence, a 3D T1 weighted sequence pre and post-Gadolinium injection, an axial dual PD-T2 weighted sequence. These patients were then split (see Table 3) between a training and testing datasets. The testing dataset was not made available to the challengers and was used to evaluate the different methods while the training dataset was provided, together with the ground truth segmentations, for challengers to train their algorithms. Images acquired on one center (3) were not part of the training dataset, with the goal of evaluating how much algorithms were dependent on the training set and sensitive to acquisition settings.

**Table 4:** Acquisition details for each sequence and each scanner for the training and testing MS patients databases.

| Scanner | Modality | Matrix | Slices | Voxel resolution (mm) |
| --- | --- | --- | --- | --- |
| GE Discovery 3T | Sagittal 3D FLAIR | 512x512 | 224 | 0.47x0.47x0.9 |
|  | Sagittal 3D T1 | 512x512 | 248 | 0.47x0.47x0.6 |
|  | Axial 2D PD-T2 | 512x512 | From 28 to 44 | 0.43x0.43x3<br>Gap: 0.5 |
| Philips Ingenia 3T | Sagittal 3D FLAIR | 336x336 | 261 | 0.74x0.74x0.7 |
|  | Sagittal 3D T1 | 336x336 | 200 | 0.74x0.74x0.85 |
|  | Axial 2D PD-T2 | 512x512 | 46 | 0.45x0.45x3 |
| Siemens Aera 1.5T | Sagittal 3D FLAIR | 256x224 | 128 | 1.03x1.03x1.25 |
|  | Sagittal 3D T1 | 256x256 | 176 | 1.08x1.08x0.9 |
|  | Axial 2D PD-T2 | 320x320 | 25 | 0.72x0.72x4<br>Gap: 1.2 |
| Siemens Verio 3T | Sagittal 3D FLAIR | 512x512 | 144 | 0.5x0.5x1.1 |
|  | Sagittal 3D T1 | 256x256 | 176 | 1x1x1 |
|  | Axial 2D PD-T2 | 240x320 | 44 | 0.69x0.69x3 |

For each patient, the challenge data includes raw datasets, and preprocessed datasets where the following steps were performed:

- Denoising of each modality using the non local means algorithm [36]
- Rigid registration of each modality on the FLAIR image [37]
- Brain extraction (skull stripping) using the volBrain platform [38], from the T1-w image and applied to other modalities
- Bias field correction of each modality using the N4 algorithm [39]

### 4.2. MS Lesions Ground Truth

Based on manual segmentation on our MS database, we first aimed at getting ground truth segmentations of multiple sclerosis lesions. This task is difficult and variability exists between experts depending on various factors, even when they follow common protocol, depending on many factors (image quality, training, modalities…). We chose to build for this challenge an unprecendented set of seven manual delineations for each patient. These delineations were performed manually on the 3D FLAIR image with control on the T2 weighted image. Each manual segmentation was performed by a trained junior expert, validated and corrected under the supervision of senior radiologists with a long experience in multiple sclerosis. More specifically, a first meeting between senior radiologists and workshop organizers of each site took place to determine the segmentation strategy and adopt a common tool (ITK-Snap) to perform manual segmentation. Junior radiologists were then recruited on each site and trained by the expert radiologists on a separate training set and when their agreement was above a threshold of 80%, they were allowed to delineate the 53 patient cases. Each case was segmented in isolation of the other cases to limit possible bias. Segmentation experts were split between the three sites which provided the patient images: 4 in Lyon, 2 in Rennes and one in Bordeaux.

MS lesions segmentation is known to be expert- and center-dependent, which can lead to relatively large discrepancies between individual manual segmentations. To cope with this problem, we computed for each patient a consensus segmentation by using the Logarithmic Opinion Pool Based STAPLE (LOP STAPLE) algorithm proposed by [10]. This algorithm computes iteratively, using an Expectation-Maximization approach, a consensus segmentation based on penalties for individual deviations from agreement between manual experts segmentations. This algorithm has several advantages: it is robust to differences between manual expert segmentations, and it allows the computation of agreement scores with respect to the consensus segmentation considered then as ground truth.

### 4.3. Computing Architecture for Automatic MS Lesions Segmentation Evaluation

One of the critical aspects in performing an independent challenge and benchmarking of medical image processing solutions is to provide a unified infrastructure able to:

- anonymize and upload the training and testing data in a single place that all participants can access though the Web
- integrate and execute the image processing algorithms through a web-based portal where all algorithms are executed on the test dataset in identical conditions
- host the processed images and make them available to the participants
- provide a cloud-based integrated solution with interoperable distributed resource management systems

Most of the past and existing challenges in the field of image processing were able to provide part of these solutions but none of them was able to provide a computing solution able to perform all of these tasks seamlessly.

We used the France Life Imaging (FLI) - Information Analysis and Management (IAM) (FLI-IAM in short) computing infrastructure for this challenge. FLI is a national infrastructure, which aims to coordinate and harmonize the network of resources on in-vivo imaging in France. Its IAM node represents the computing node of France Life Imaging (**https://www.francelifeimaging.fr/en/about/noeuds/iam/**). This architecture allows (see Figure 7) the storage and management of preclinical and clinical *in vivo* imaging data and offers services of images processing and analysis.

**Figure 7:**
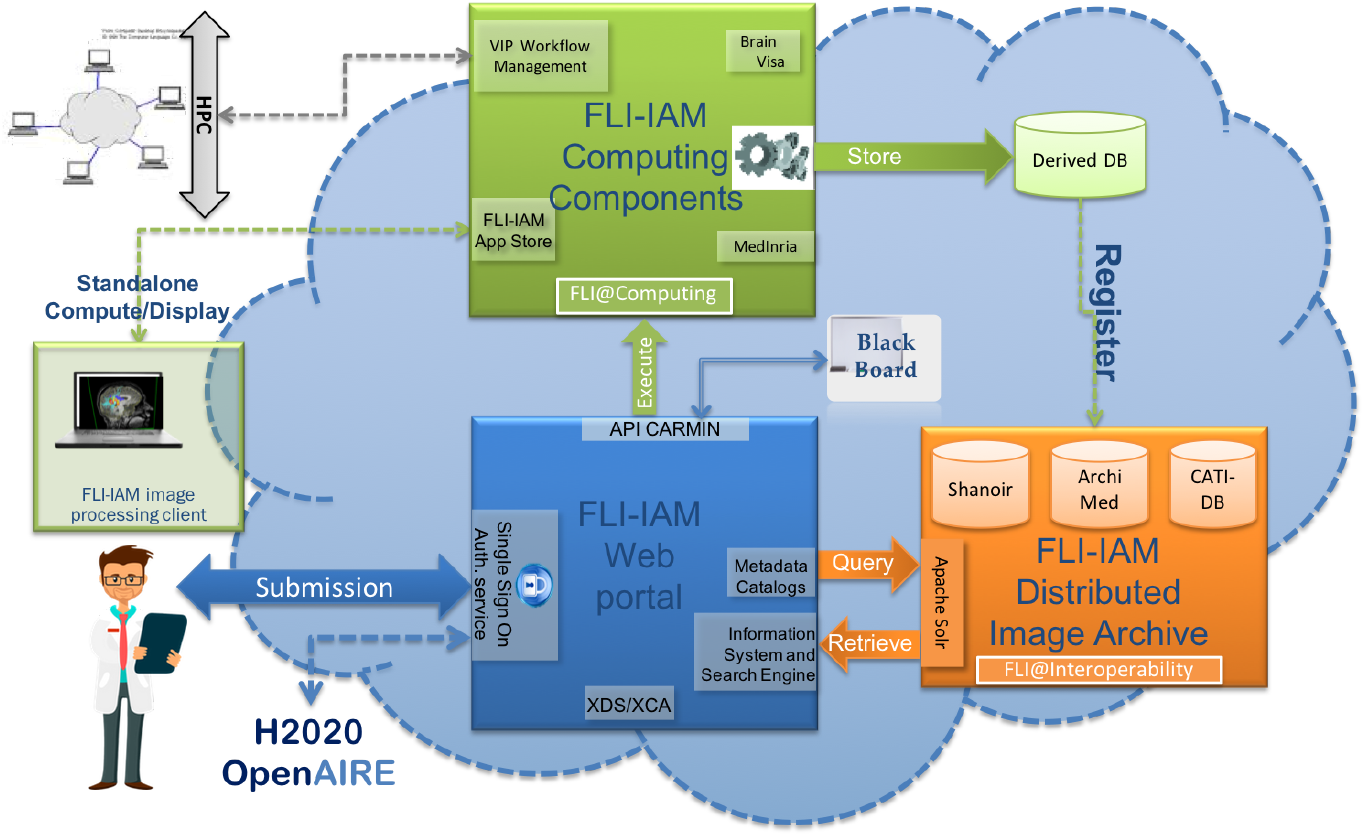
FLI-IAM architecture.

FLI-IAM is based on existing, technologically ready software solutions, coming from multiple research teams over France. It proposes a web portal (blue box in Fig. 7) to unify the access to all resources and tools and provides multiple solutions for storage and computation on medical images.

To operate this challenge, three components of the entire FLI-IAM portfolio have been used:

- Web portal
- Shanoir (SHAring NeurOImaging Resources) for the database [40]
- VIP (Virtual Imaging Platform) for the computing platform [41]

In addition to these three major components, additional tools/services have been developed to provide the required level of interoperability and to finally integrate all components into one unique workflow.

The web portal has been used as a communication platform with all challengers. All information concerning the challenge has been distributed there, e.g. the organizational aspects, dataset descriptions, evaluation details, etc. Challengers had to subscribe on the portal to participate in the challenge.

Shanoir (SHAring NeurOImaging Resources) served as central database for all datasets necessary for the challengers, all their processed results and challenger’s scores. Shanoir is an open source neuro-informatics platform designed to share, archive, search and visualize neuroimaging data. It provides a user-friendly secure web access and offers an intuitive workflow to facilitate the collection and retrieval of neuroimaging data from multiple sources. Shanoir comes along many features such as anonymization of data, support for multi-center clinical studies on subjects or group of subjects.

VIP (Virtual Imaging Platform) provided all necessary resources for the integration and the execution of all challenger processing pipelines. The pipelines were provided by challengers as Docker containers and were integrated into VIP using the Boutiques application repository. Boutiques relies on Linux containers to solve the problem of application installation in a lightweight manner and it uses a versatile JSON format to describe command line tools. VIP also ensured the execution of the challenger pipelines (and the subsequent segmentation performance analysis) on the computing resources available for the challenge.

Figure 8 gives an overview of the integration level between database and computing platform and describes the workflow that was set up for the hosting of the challenge.

**Figure 8:**
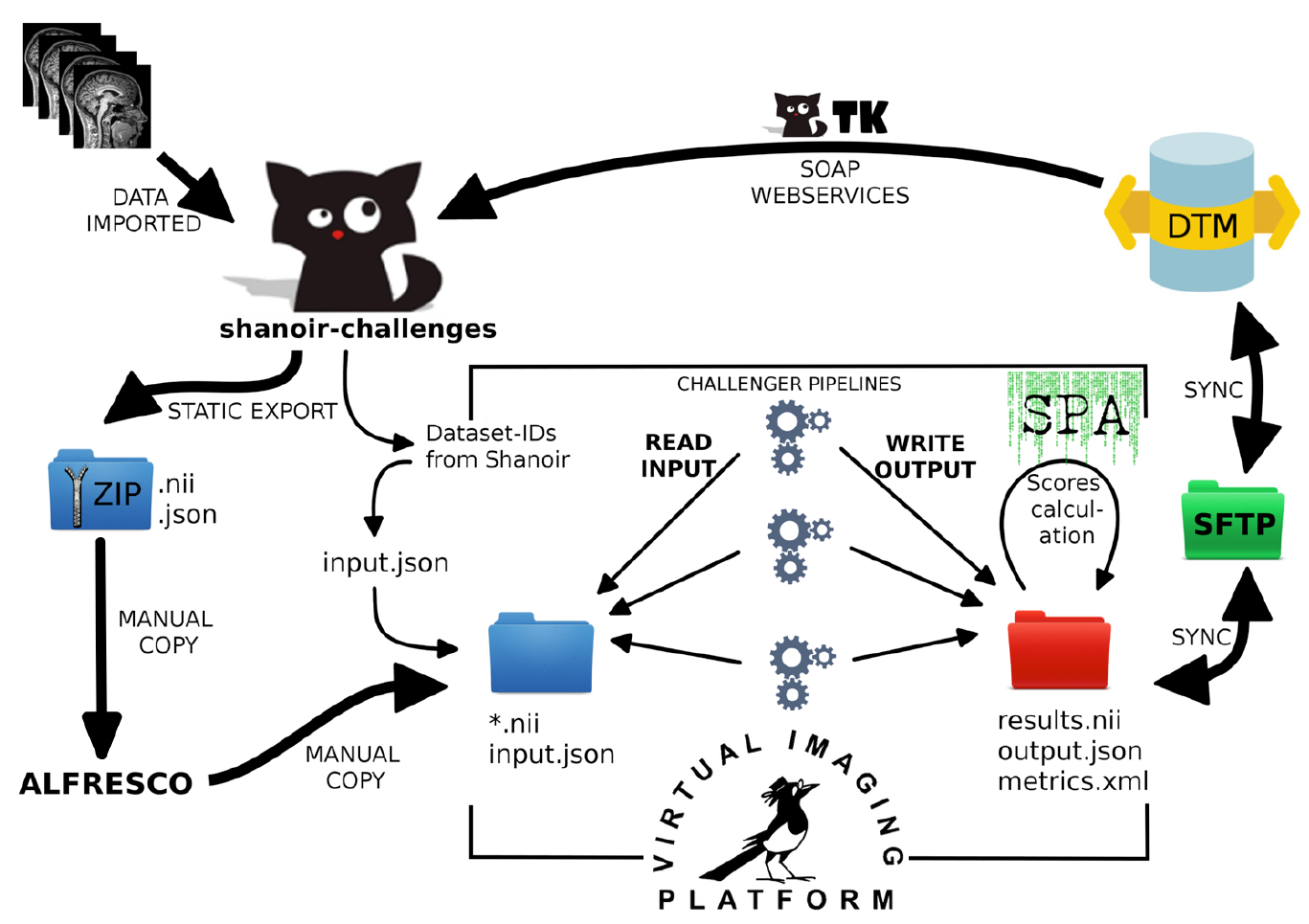
Workflow for database and computing platform integration.

After the preparation of the challenge data, all source datasets were imported into Shanoir. The training data were shared with the challengers using the portal and its file download feature. The testing data was processed by VIP using a static export folder exported from the database - containing all necessary metadata from the database to import results back into the database and attach them to the source dataset for each challenger.

After the algorithms produced their results, the segmentation performance analysis (including all measures described in the next section) was run to compare challengers results with the consensus ground truth and calculate the scores. The continuously running DataTransferModule has been connected to the results folder of VIP to automatically import back the result datasets into Shanoir. All result datasets and their corresponding scores have been made available in Shanoir for challengers. The challenge organization team could then easily access the scores summary within Shanoir.

### 4.4. Challenge Evaluation Strategy and Metrics

Using the computing platform, challengers were asked to provide algorithms delineating lesions in the FLAIR reference space. This was asked to match the space in which the manual delineations were carried out and therefore avoid any unwanted discrepancies due to interpolation of the challengers’ results. To evaluate these results, we have implemented a large set of evaluation measures for the challenge with the goal of evaluating the different aspects evaluated by clinicians when looking at MS patient images. For this reason, we have separated the evaluation into two major categories of evaluation metrics:

- Segmentation evaluation: does the algorithm provide a precise delineation of each lesion?
- Lesion detection evaluation: does the algorithm find all lesions in the image independently of its precise delineation?

Each of these categories may contain several metrics that characterize differently the segmentation quality. We describe in more details each of the chosen metrics for the challenge in the following sections. All evaluation algorithms used for this paper are available open-source as part of the Anima software (**http://github.com/Inria-Visages/Anima-Public**). Although not presented in this article (but available as part of supplementary material **(http://dx.doi.org/10.5281/zenodo.1307653)**, we remind the strategy that was used at the challenge to rank, for each of these metrics, the different methods on all patients:

- For each patient, compute the selected metric for each algorithm by comparing it to the ground truth previously computed
- For each patient, rank the algorithms according to the selected metric (from 1: best performing to *N*: worst performing)
- Compute for each algorithm its average ranking over all patients evaluated. This average rank is used for the final ranking of the methods

We had selected this approach instead of simply averaging the metric scores for each algorithm to avoid a bias of some methods that would get a few very good metric scores that would not represent their true behavior. This approach instead considers as the best method the one that ranks the best on average for all patients evaluated, thereby discarding this bias problem. Instead in this work, we focus more on the graphical analysis of the cluster analysis of the algorithms with respect to the experts who delineated the structures. To this end, we performed a multi-parametric analysis of the results. For each couple of metrics presented in the following (average surface distance, Dice score and *F*_1_ score), we computed a 2D scatter representation of the average results on all testing patients of each of the teams and of the experts. Since different clusters of results quality may be outlined by such graphs, we then ran for each combination of metrics a clustering into three groups of the average performance of the teams and experts. For this clustering to be precise enough however, we need to account for the variance around the average points. We have therefore chosen to perform a spectral clustering [42], considering each point of the 2D graph not as a mean but as a multivariate Gaussian, using a distance between multivariate Gaussians as expressed in [43], thus accounting for the covariance in the individual scores.

#### 4.4.1. Segmentation evaluation

The first category of evaluation metrics is also the most known in the literature and concerns segmentation evaluation, i.e. are the contours of the lesions precisely delineated compared to the ground truth. In this group, we distinguish two sub-categories, each quantifying the precision of lesions delineation: overlap-based and surface-based metrics. In the following, we will consider two binary images representing respectively the lesions consensus (i.e. the ground truth): *G*, and the evaluated segmentation (i.e. one algorithm segmentation result): *A*, both illustrated in Fig. 9.

**Figure 9:**
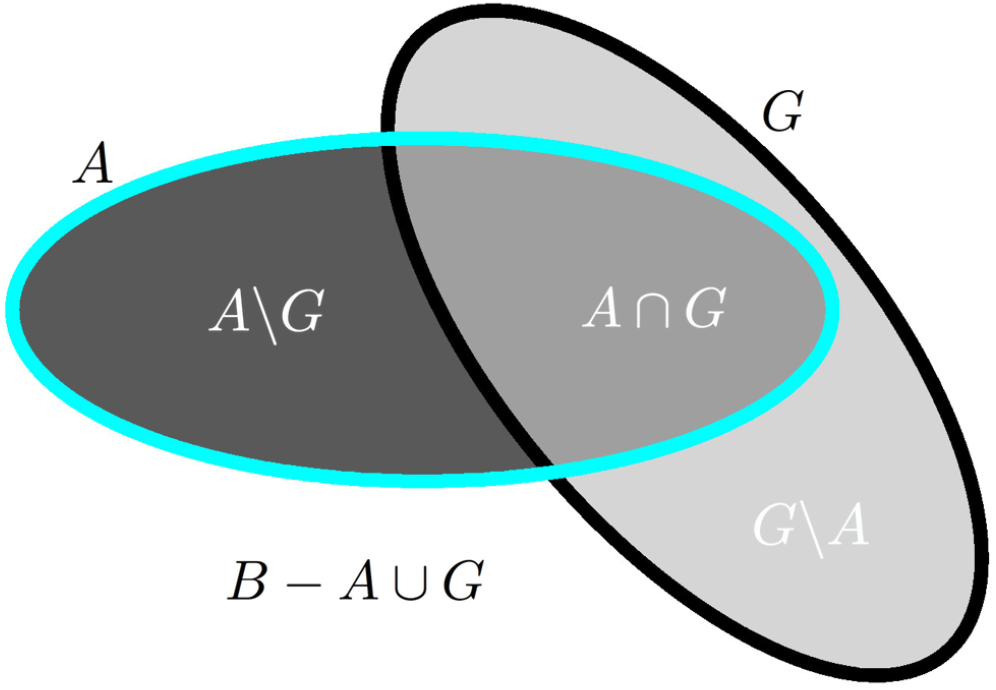
Illustration of overlap-based segmentation evaluation: quantities used for measures computation. *A* denotes the evaluated segmentation, *G* the ground truth, and *B* the image domain.

*Overlap metrics*. These measures consider the voxel-based overlap of A and G based on the quantities illustrated in Fig. 9. Among those measures, we use the following ones:

- Dice score [44]: 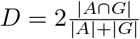
- Positive predictive value: 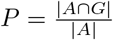
- Sensitivity: 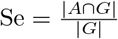
- Specificity: 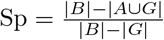

where *A* ∪ *G* is computed from other quantities: *A* ∪ *G* = *A* ∩ *G* + *A*\*G* + *G*\*A*. For all formulas in this section, the notation |.| denotes taking the cardinal of a set of voxels, e.g. |*A* ∩ *G*| denotes the number of voxels in that set. As a final remark for this category, the choice of the size of image *B* is quite important as it will influence specificity. A too large region for *B* could indeed lead all specificity values to be very close to 1 by construction and therefore make them difficult to compare. We therefore chose for the challenge to compute *B* as the union of all available segmentations for a patient (automatic and manual), dilated three times by a 6-connectivity kernel.

Each overlap-based metric varies between 0 and 1, 1 being a perfect result and 0 the worst result. Each measure is however sensitive to a different phenomenon in the quality of segmentations: positive predictive value and specificity are influenced by false positives and are therefore sensitive to overly large segmentations; sensitivity is influenced by false negatives and is thus sensitive to overly small segmentations. Finally, the Dice score is a composite measure attempting to summarize all influences into a single scalar measure.

*Surface metric*. In addition to overlap-based metrics, we have computed the average symmetric surface distance, also used in MICCAI 2008 challenge on MS lesions segmentation organized by [7]. Instead of using voxel-based overlaps, this measure uses contours extracted from the two input segmentations *A* and *G*, denoted respectively *A_S_* and *G_S_*. This distance is expressed as the following sum:

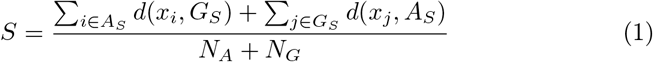

where *d* denotes the minimal Euclidean distance between a point of one surface and the other surface, *N_A_* and *N_G_* denote the number of points of each surface.

#### 4.4.2. Detection evaluation

As mentioned in the introduction, evaluation of the detection of lesions is as crucial, if not even more, as segmentation precision as the number of lesions is used for MS diagnosis. We wanted to evaluate in this category how many lesions have been (in)correctly detected, independently of the precision of their contours.

*Defining lesion detection*. This whole category of measures relies on identifying individual lesions in the ground truth *G* and evaluated segmentations. For this task, we first compute the connected components of G and A (with a 18- connectivity kernel) and remove all lesions that are smaller in size than 3 mm^3^. We therefore get label images 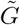 and *Ã* where each label denotes a specific lesion.

From these two labeled images, two quantities are computed that will be used to characterize the detection power of an algorithm:

- TP*_G_*: the number of lesions among the *M* lesions in the ground truth 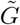 that are correctly detected by *Ã*
- TP*_A_*: the number of lesions among the *N* lesions in the automatic segmentation *Ã* that are correctly detected by 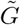

Let us consider only the case of TP_*G*_, TP*_A_* being computed with the same procedure but reverting the roles of *Ã* and 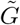. We first construct the joint histogram *H* of *Ã* and 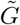 where *H_i,j_* corresponds to the number of voxels having label *i* ∈ {0 … *M*} in 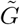 and label *j* ∈ {0 … *N*} in *Ã*. We consider a lesion *j* in 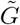 (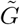_*j*_) to be detected if it respects the following rules:

- The lesion 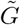*j* is overlapped at least at a rate of α% by lesions of *Ã*
- Lesions of *Ã* that contribute the most to the detection of 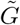*j* (summing up to γ% of the total overlap) do not go outside of 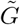*j* by more than *β*%.

While the first condition ensures that the lesion to be detected is sufficiently overlapped, the second condition ensures that the detection is not due to an overly large segmentation in *Ã* that would overlap many lesions in 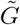 by chance. These two conditions are implemented in Algorithm 1.

##### Algorithm 1 TP_*G*_ computation algorithm

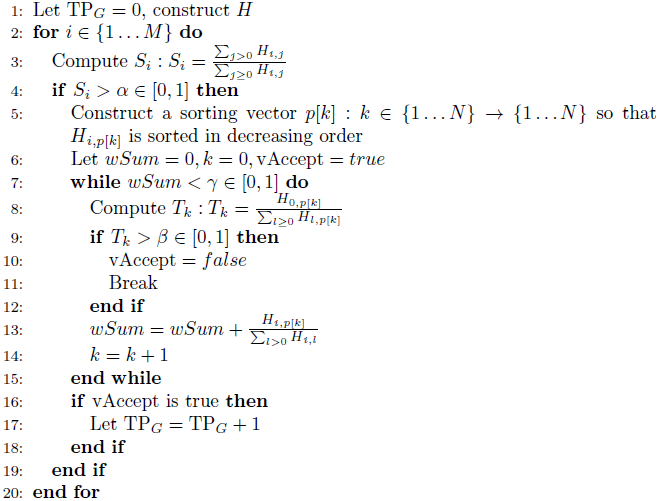

For the challenge, we used this algorithm with values heuristically defined on several independent tests to give meaningful values for TP_*G*_ and TP_A_: *α* = 10%, γ = 65%, *β* = 70%.

*Detection metrics*. From the number of lesions *M* and *N* respectively in 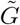 and *Ã*, and the numbers computed above (TP_*G*_ and TP_*A*_), the following detection metrics are computed, named after their similarity to overlap-based metrics:

- Lesion sensitivity, i.e. the proportion of detected lesions in 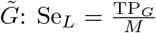
- Lesion positive predictive value, i.e. the proportion of true positive lesions inside 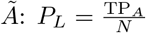

In addition to these two metrics, we have computed a summary metric to get, like the Dice score for segmentation metrics, a one-glance idea of the detection performance of a given method (0 meaning worst performance and 1 meaning perfect detection performance). This summary metric, the *F*_1_ score, considers both lesion sensitivity and positive predictive value to compute the score. It is defined as follows:

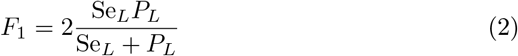

## Acknowledgments

This work was partly funded by France Life Imaging (grant ANR-11-INBS-0006 from the French “Investissements d’Avenir” program) for funding and sponsoring the challenge. This work has also been partly supported by a grant (OFSEP) provided by the French State and handled by the “Agence nationale de la recherche”, within the framework of the “Investissements d’Avenir” program, under the reference ANR-10-COHO-002. We also thank the French national cohort OFSEP (a French “Investissements d’Avenir” program), and particularly the imaging group inside this cohort consortium for their constant support, fruitful discussions on the challenge and providing the MR images.

## Additional Information

Authors of this article contributed to the article in the following manner:

- Jérémy Beaumont, Olivier Commowick, Christian Barillot, Senan Doyle, Michel Dojat, Florence Forbes, Jesse Knight, April Khademi, Amirreza Mahbod, Chunliang Wang, Richard McKinley, Franca Wagner, John Muschelli, Elizabeth Sweeney, Eloy Roura, Xavier Lladó, Michel M. Santos, Wellington P. Santos, Abel G. Silva-Filho, Xavier Tomas-Fernandez, Simon K. Warfield, Hélène Urien, Isabelle Bloch, Sergi Valverde, Mariano Cabezas, Francisco Javier Vera-Olmos and Norberto Malpica designed the challengers’ repsective algorithms, participated to the challenge, participated in the writing and proof-reading of the evaluated teams description in particular, and of the proof-reading of the whole article.
- Michaël Kain, Baptiste Laurent, Florent Leray, Mathieu Simon, Sorina Camarasu Pop, Pascal Girard, Frédéric Cervenansky, Tristan Glatard, Olivier Commowick, Christian Barillot and Michel Dojat participated in the setup of the platform and running of the experiments on the France Life Imaging platform, the writing and proof-reading of the pipeline processing description and results in particular, and of the proof-reading of the whole article.
- Olivier Commowick, Audrey Istace, Florent Leray, Baptiste Laurent, Roxana Améli, Jean-Christophe Ferré, Anne Kerbrat, Thomas Tourdias, Sandra Vukusic, Gilles Edan and François Cotton participated in the constitution of the evaluation database (selection of the patients, expert guidance on the delineation of the lesions), in the analysis of results, and in the writing of the corresponding sections of the paper in particular. They also participated in the proof-reading of the whole article.
- Olivier Commowick, Charles Guttmann, Frédéric Cervenansky, Martin Styner, Simon K. Warfield, François Cotton and Christian Barillot participated in all aspects of the challenge organization, design of the results evaluation experiments and metrics, writing and proof-reading of all sections of the paper.

The authors of this article have no competing interests as defined by Nature Research, or other interests that might be perceived to influence the results and/or discussion reported in this paper.

